# Sodium arsenite requires redox sensitization to cause developmental dopaminergic neurotoxicity in *Caenorhabditis elegans*

**DOI:** 10.64898/2026.10.30.763158

**Authors:** Javier Huayta, Wolmer Espinoza Tapia, Jay Jhaver, Joel N. Meyer

## Abstract

Environmental pollutants can disrupt mitochondrial function through many pathways and mitochondrial dysfunction is often linked to neurotoxicity, particularly of dopaminergic neurons. However, there are many mechanisms of mitochondrial toxicity. Remarkably, despite the centrality of the Krebs cycle to mitochondrial bioenergetics, there is very limited evidence regarding whether pollutants that inhibit the Krebs cycle can cause neurotoxicity. We exposed *Caenorhabditis elegans* through its whole development to Krebs cycle inhibitors sodium arsenite and fluoroacetic acid, and evaluated neuronal morphology and function as well as dopaminergic ATP levels and redox state. Both chemicals impaired growth and reproduction, but neither caused morphological damage nor functional deficits in dopaminergic, glutamatergic, or cholinergic neurons at concentrations causing mild organismal toxicity. Sodium arsenite, but not fluoroacetic acid, decreased dopaminergic ATP levels; neither caused significant changes in mitochondrial redox state. However, developmental exposure to sodium arsenite caused dopaminergic neurotoxicity when mitochondrial quality control or redox stress response was compromised by loss of *pink-1*/PINK1 or *sod-2*/SOD2, by co-exposure to rotenone, antimycin A, or buthionine sulfoximine, or secondary challenge to 6-hydroxydopamine. These conditions also led to dopamine-associated behavioral deficits. Thus, developmental exposure to sodium arsenite required an additional mitochondrial or redox stress challenge to cause dopaminergic neurotoxicity, and we did not find evidence that inhibition of the Krebs cycle broadly defined causes developmental neurotoxicity.

## 1. Introduction

Arsenic exposure remains an important environmental health concern. Inorganic arsenic occurs naturally in groundwater and can also enter drinking water, food, and air through industrial and agricultural activities [1, 2]. Chronic exposure to arsenic is associated with cancer, cardiovascular and metabolic disease, and adverse developmental outcomes [3, 4], while developmental and early-life exposure has been linked to persistent health effects that may emerge long after the original exposure [5–7]. Neurological and cognitive effects of arsenic exposure have also been reported, particularly following developmental exposure, but the cellular mechanisms that connect arsenic exposure to neurotoxicity are not completely understood [8, 9]. Moreover, mitochondria are a major target of arsenic toxicity [10–13], while also being important for the etiology of Parkinson’s disease (PD) and other neurodegenerative disorders [14, 15]. However, mitochondrial dysfunction comprises many mechanistically distinct pathways, such as inhibition of all the respiratory chain complexes, impaired tricarboxylic acid (Krebs) cycle activity, redox cycling, uncoupling of ATP production from oxygen consumption, and mitochondrial DNA damage. These mechanisms can produce different outcomes after pollutant exposure; for example, a mitochondrial toxicant may reduce ATP availability without producing a shift towards an oxidative environment, while inhibition of particular electron transport chain sites can generate reactive oxygen species leading to neurotoxicity [16–18]. Therefore, treating mitochondrial dysfunction as a single mode of action may obscure the mechanisms most relevant to the onset of neurotoxicity and neurodegenerative disease.

The Krebs cycle is central to supply reducing equivalents to oxidative phosphorylation and intermediates for biosynthesis [19, 20], making it a potentially critical target for environmental toxicants and disruption of neuronal energy metabolism. Sodium arsenite can inhibit many enzymes, including several involved in pyruvate metabolism and Krebs cycle function and, in *C. elegans*, decreases ATP-linked respiration and spare respiratory capacity while increasing proton leak and reliance on glycolytic metabolism [12], with distinct genetic backgrounds influencing its effects [21, 22]. The pesticide and chemical metabolite fluoroacetic acid provides a mechanistically distinct and more specific approach to Krebs cycle inhibition; it is converted to fluorocitrate, which inhibits aconitase, an enzyme shared by the Krebs and glyoxylate cycles [23, 24]. Moreover, fluoroacetic acid toxicity in mice is characterized by seizures [25], with fluoroacetate injection causing memory loss in chicks [26, 27]. In people, reports of poisoning with fluoroacetate show a link to long-term cerebellar ataxia [28], and seizures and altered consciousness [29]. These cases point to potential and persistent effects in the central nervous system. Additionally, previous studies show that sodium arsenite effects in *C. elegans* depend strongly on life stage, genetic background, and toxicological endpoints quantified [12, 13].

Finally, chronic millimolar arsenite exposure has been reported to induce dopaminergic neurodegeneration and behavioral impairment, accompanied by activation of oxidative stress responses [24]. These outcomes showcase the importance of assessing the potential neurotoxic effects of arsenic because of its ubiquitous environmental presence and capacity to disrupt mitochondrial function.

In this study, we used *C. elegans* to determine whether developmental inhibition of Krebs cycle by exposure to sodium arsenite and fluoroacetic acid is sufficient to cause dopaminergic, glutamatergic, or cholinergic neurotoxicity *in vivo*. We exposed worms from the embryo stage to the L4 larval stage to ensure that our endpoints were quantified after all somatic cells of *C. elegans* were developed. We then evaluated neuronal morphological damage and loss of function. We assessed mitochondrial redox and bioenergetic states and tested whether genetic or chemical interventions that reduce redox stress response or bioenergetic capacity enabled neurotoxicity under concentrations causing mild organismal toxicity. We found that neither sodium arsenite nor fluoroacetic acid by themselves caused neurotoxicity or neuron functional deficit under these conditions. However, sodium arsenite caused dopaminergic neurotoxicity when redox stress response was compromised by secondary chemical or genetic challenge. Our results indicate that developmental sodium arsenite exposure can create a state of dopaminergic neuron vulnerability in which additional mitochondrial or oxidative stress is required to produce neurotoxicity, but do not support that Krebs cycle inhibition *per se* causes developmental neurotoxicity.

## 2. Materials and Methods

### 2.1. C. elegans maintenance

2.1.1 *C. elegans strains*

Strains BY200 (*vtIs1*[p*dat-1*::GFP]), DA1240 (*adIs1240*[*eat-4*::sGFP+lin-15(+)]), LX929 (*vsIs48*[*unc-17*::GFP]), CB1112 (*cat-2*(e1112) II), CB1141 (*cat-4*(e1141) V), CB75 (*mec-2*(e75) X), CB1339 (*mec-4*(e1339) X), NM1657 (*unc-10*(md1117) X), VC223 (*tom-1*(ok285) I), UA227 (*pdr-1*(tm598); *vtIs1*[p*dat-1*::GFP]), UA226 (*pink-1*(tm1779); *vtIs1*[p*dat-1*::GFP]), ERS1 (*eraIs1*[*dat-1*p::mCherry, *dat-1*p::hSNCA::Venus]), ERS44 (*eraIs1*[*dat-1*p::mCherry, *dat-1*p::hSNCA::Venus]; *pdr-1*(gk448)), ERS49 (*eraIs1*[*dat-1*p::mCherry, *dat-1*p::hSNCA::Venus]; *pink-1*(tm1779)), PHX2867 (p*dat-1*::MLS::roGFP), PHX2923 (p*dat-1*::PercevalHR), JV2 (*jrIs2*[*rpl-17*p::Grx1-roGFP2 + *unc-119*(+)]), BY200 x GA184 (*sod-2*(gk257; *vtIs1*[p*dat-1*::GFP]) I), BY200 x GA186 (*sod-3*(tm760) X; *vtIs1*[p*dat-1*::GFP]), GR2245 (*skn-1*(mg570) IV); QV225 (*skn-1*(zj15) IV), *aak-1*(tm1944) III, and TG38 (*aak-2*(gt33) X) were maintained at 20 °C on K-agar plates [30], seeded with OP50 *E. coli*. Strain BY200 was a gift from Michael Aschner, strains ERS1, ERS44, and ERS49 were a gift from Roman Vozdek, and other strains were obtained from the Caenorhabditis Genetics Center and the National BioResource Project (<u>Table A1</u>).

#### 2.1.2 Generation of age-synchronized populations

A non-starved population of day 1 adults was collected in a 15 mL conical tube from K-agar plates by washing twice with K-medium [31]. Worms were gravity-settled for 3 minutes, and the supernatant discarded. Worms were treated with 5 mL of K-medium containing final concentrations of 0.4 N sodium hydroxide and 20% v/v sodium hypochlorite, and the conical tube put on a rocking shaker for 8 minutes. The bleaching reaction was quenched by raising the volume to 15 mL with K-medium, centrifuged at 2200 RCF for 2 minutes, and the supernatant discarded. This washing step was repeated two more times. Embryos were counted by transferring 10 µL of the remaining liquid to a glass slide and observed on a Leica MZ75 stereomicroscope. K-medium was added or subtracted until reaching an approximate concentration of 10 embryos per 1 µL. At this point, embryos were ready for developmental exposure experiments.

#### 2.1.3 Generation of crossed strains

Approximately 40-50 L4-stage worms of strain BY200 were exposed to 33 °C for 4 hours on K-agar plates to generate a male-enriched population. Worms were then transferred to 15 °C overnight and moved to 20 °C the next day. After 3-4 days, the offspring of heat-shocked worms were screened for males and transferred to new plates containing three hermaphrodites and 10-12 males each. This step was necessary to increase the number of available males in the next generation. After 3-4 days, plates were prepared containing three hermaphrodites of either of the GR2245, QV225, tm1944 or TG38 strains and 10-12 BY200 second-generation males. After 3-4 days, individual offspring worms with visible BY200 dopaminergic neuron fluorescence were transferred to fresh plates. After 3-4 days, putative homozygous plates were screened for visible BY200 dopaminergic neuron fluorescence. Putative homozygous crosses were verified by phenotyping for strain QV225, and by PCR genotyping for strains GR2245, tm1944, and TG38 (<u>Table A2</u>).

### 2.2. Chemical exposure design

#### 2.2.1 Developmental exposure

Approximately 100 embryos in 10 µL of K-medium from the bleaching treatment were transferred to each well of a 24-well plate and the volume increased to 500 µL per well of complete K-medium (0.5 mL of 10 mg/mL cholesterol, 3 mL of 1 M CaCl_2_, 3 mL of 1 M MgSO_4_ per 1 L K-medium) containing HB101 *E. coli* bacterial food at a final concentration of 5 mg/mL, and either 10 µL of sodium arsenite (Sigma-Aldrich) or fluoroacetic acid (Sigma-Aldrich) stock solutions. The 24-well plate was put on an orbital shaker at 20 °C and 120 RPM for 52-54 hours to allow worms to reach the L4 larval stage. Afterward, liquid from each well was collected in separate 15 mL conical tubes, the total volume raised to 15 mL with K-medium, and the worms were allowed to gravity-settle. After 3 minutes, the supernatant was discarded, and this washing step was repeated two more times. At this point worms were ready for developmental endpoint measurement.

#### 2.2.2 Determination of experimental concentrations

Embryos were treated as described in the preceding paragraph. Final concentrations tested were 0, 50, 250, 500, 750, 1000, 1500 and 2000 µM sodium arsenite (Sigma-Aldrich), or 0, 50, 100, 250, 500, 750, 1000 and 1500 µM fluoroacetic acid (Sigma-Aldrich). After 52-54 hours, worms were transferred to unseeded K-agar plates, allowing the worms to disperse on the plate for 5 minutes. Plates containing worms were mounted and imaged using a Keyence BZ-X710 microscope using a Nikon 4X objective. Images were analyzed using the WormSizer add-in in ImageJ following their published protocol to measure length and volume of 50-100 worms per treatment [32]. These length and volume values were used to construct a dose response curve using the AAT Bioquest EC_50_ Calculator to determine EC_1_, EC_10_, EC_25_, and EC_30_ concentrations [33].

### 2.3. Quantification of viable offspring

Approximately 100 embryos in 10 µL of K-medium from the bleaching treatment were transferred to each well of a 24-well plate and the volume increased to 500 µL per well of complete K-medium (0.5 mL of 10 mg/mL cholesterol, 3 mL of 1 M CaCl_2_, 3 mL of 1 M MgSO_4_ per 1 L K-medium) containing HB101 *E. coli* bacterial food at a final concentration of 5 mg/mL, and either 0, 250, 500, and 1000 µM sodium arsenite (Sigma-Aldrich) or 0, 50, 250, and 500 µM fluoroacetic acid (Sigma-Aldrich) final concentrations. The 24-well plate was put on an orbital shaker at 20 °C and 120 RPM for 52-54 hours to allow worms to reach the L4 larval stage.

Afterward, liquid from each well was collected in separate 15 mL conical tubes, the total volume raised to 15 mL with K-medium, and the worms were allowed to gravity-settle. After 3 minutes, the supernatant was discarded, and this washing step was repeated two more times. Worms were transferred to K-agar plates containing 1 worm per plate for a minimum of 8 worms per treatment. Worms were transferred to fresh K-agar plates every 24 hours for 5 days; only larvae but not unhatched embryos were counted every day.

### 2.4. Quantification of neurotoxicity

#### 2.4.1 Scoring of neuronal damage

After developmental chemical exposure, 20-30 worms were transferred to a 2% agarose pad on a glass slide containing a drop of 100 mM sodium azide to cause paralysis. After 3 minutes, paralyzed worms on the agarose pad were covered with a coverslip. Z-stacks of individual worms’ heads were acquired on a Keyence BZ-X710 microscope equipped with a Keyence BZ-X700E metal halide light-source, using a Nixon 40X (air) objective with 200 milliseconds of exposure, pitch of 0.5 µm, and the “high sensitivity” option selected. Maximum projections of the Z-stacks were generated based on maximum intensity, and each dendrite of the cephalic neurons was scored as previously described [34, 35]. Briefly, for dopaminergic neurons: 0 – no damage, 1 – irregular (curves), 2 – less than 5 blebs, 3 – 5 to 10 blebs, 4 – more than 10 blebs and/or breaks, 5 – breaks, 25 to 75% dendrite loss, and 6 – breaks, more than 75% dendrite loss; for glutamatergic neurons: 0 – no damage, 1 – 1 to 5 blebs, 2 – 6 to 10 blebs, and 3 – more than 10 blebs and/or presence of breaks; and for cholinergic neurons: 0 – no damage, 1 – irregular (curves), 2 – 1 to 10 blebs, and 3 – more than 10 blebs and/or presence of breaks (<u>Figure A1</u>)

#### 2.4.2 Basal slowing response

Assay plates were prepared by spreading 200 µL of HB101 *E. coli* cultured in LB Broth in the shape of a ring on K-agar plates and incubated overnight at room temperature. After developmental chemical exposure, worms were collected and transferred to a 15 mL conical tube and allowed to gravity-settle for 1-2 minutes. After discarding the supernatant, 30-40 worms were transferred to the center of either an assay plate or a plate with no bacteria, with excess liquid absorbed using a Kimwipe. After 5 minutes, a minimum of 5 videos of 20 seconds each per plate were acquired using a Keyence BZ-X710 microscope. These videos were used to count the number of body bends in 20 seconds of approximately 10 worms per plate. A body bend was counted each time the portion of the worm corresponding to the base of the pharynx reached the maximum amplitude of its crawling sinusoidal wave. Strains CB1112 and CB1141, defective for basal slowing response, were used as positive controls.

#### 2.4.3 Sensitivity to soft touch

After developmental chemical exposure, worms were collected and transferred to a 15 mL conical tube and allowed to settle for 3 minutes. After discarding the supernatant, 30-40 worms were transferred to the center of a K-agar plate with no bacteria and allowed to dry for 5 minutes. Worms were tested by stroking their heads and tails consecutively with an eyelash mounted on a Pasteur pipette under a Leica MZ75 stereomicroscope. Sensitive worms started backwards movement after their heads were touched and stopped or started forward locomotion after their tails were touched. This process was repeated 5 times per worm for a total of 20 worms per replicate, counting the number of times that a worm showed response to soft touch. Worms that did not start backwards motion after the initial head touch three consecutive times were also counted as not responsive for that try. Strains CB75 and CB1339, defective for sensitivity to soft touch, were used as positive controls.

#### 2.4.4 Aldicarb-induced paralysis

Assay plates containing 1 mM aldicarb (Sigma-Aldrich) were prepared one day before the assay and stored at 4 °C. Plates were allowed to reach room temperature before starting the assay. A drop of 2 µL of OP50 *E. coli* was added to the center of the plate and allowed to dry for 5 minutes. After developmental chemical exposure, worms were collected and transferred to a 15 mL conical tube and allowed to gravity-settle for 3 minutes. After discarding the supernatant, 25-30 worms were transferred to the bacterial spot on the plate and paralysis was assessed every 30 minutes for 6 hours. A paralyzed worm was defined as not showing movement after prodding with an aluminum wire. Strains NM1657 and VC223, resistant and hypersensitive to aldicarb paralysis respectively, were used as positive controls.

### 2.5. Quantification of mitochondrial redox and ATP states

Strains PHX2867 and PHX2923, expressing mitochondrial-localized dopaminergic roGFP and PercevalHR respectively under the *dat-1* promoter [36], were used to quantify the glutathione redox tone and ATP:ADP ratio in the cell bodies of the cephalic dopaminergic neurons. Strain JV2, ubiquitously expressing the roGFP2 sensor for estimation of GSSG/2GSH ratio, was used to quantify redox tone in the whole body of worms. After developmental chemical exposure, worms were collected and transferred to a 15 mL conical tube and allowed to gravity-settle for 3 minutes. After discarding the supernatant, 20-30 worms were transferred to an 8% agarose pad on top of a glass slide and covered with a coverslip. The glass slide was mounted on a Keyence BZ-X710 microscope equipped with a Keyence BZ-X700E metal halide light-source. For strains PHX2867 and PHX2923, two images per head of an individual worm, focused on the cell bodies of the cephalic dopaminergic neurons, were acquired using a Nikon 40X (air) objective, 150 and 70 milliseconds of exposure for the GFP and roGFP channels respectively, and with the “high sensitivity” option selected. The first image was acquired using a 470ex/520em filter, and the second image with a 405ex/520em filter. For strain JV2, the imaging process was the same, except that both images were captured for the whole body of the worm using a Nikon 10X objective. A custom-made MATLAB script (available from Duke Libraries Repository [37]) was used to perform feature-based segmentation of the cell bodies in each image, background subtraction, and to determine their mean intensity values. The script then calculated the mean intensity ratios of 405/470 excitation for mitochondrial roGFP as oxidized to reduced glutathione, and the mean intensity ratios of 470/405 excitation for PercevalHR as ATP to ADP [17]. A modified version of the script was used for quantification of roGFP ratios for the JV2 strain, to accommodate the segmentation of the whole body instead of only the cephalic dopaminergic cell bodies.

### 2.6. Secondary chemical co-exposure

#### 2.6.1 Chemical sensitization

Developmental exposure was performed as described for single chemicals, with the following modifications. Approximately 100 embryos in 10 µL of K-medium from the bleaching treatment were transferred to each well of a 24-well plate and the volume increased to 500 µL per well of complete K-medium (0.5 mL of 10 mg/mL cholesterol, 3 mL of 1 M CaCl_2_, 3 mL of 1 M MgSO_4_ per 1 L K-medium) containing HB101 *E. coli* bacterial food at a final concentration of 5 mg/mL For each replicate, one well contained no additional chemicals to serve as control, one well contained only the additional chemical tested for sensitization, one well contained the EC_10_ or EC_25_ sodium arsenite (Sigma-Aldrich) concentrations, and one well contained the EC_10_ or EC_25_ sodium arsenite concentrations and the sensitization chemical. Final concentrations of the sensitization chemicals were 1.0 µM rotenone (Sigma-Aldrich), 0.1 µM antimycin A (Sigma-Aldrich), or 500 µM buthionine sulfoximine (Sigma-Aldrich). All wells also contained a final concentration of 1% v/v DMSO as vehicle. The 24-well plate was put on an orbital shaker at 20 °C and 120 RPM for 52-54 hours to allow worms to reach the L4 larval stage. Afterward, liquid from each well was collected in separate 15 mL conical tubes, the total volume raised to 15 mL with K-medium, and the worms were allowed to settle. After 3 minutes, the supernatant was discarded, and this washing step was repeated two more times. At this point worms were ready for developmental endpoint measurement.

#### 2.6.2 6-hydroxydopamine exposure

After developmental chemical exposure, worms were collected and washed as described for single chemicals. These worms were then transferred to 24-well plates and well volumes were raised to a total of 500 µL per well of complete K-medium (0.5 mL of 10 mg/mL cholesterol, 3 mL of 1 M CaCl_2_, 3 mL of 1 M MgSO_4_ per 1 L K-medium) containing 25 or 50 mM 6-hydroxydopamine (6-OHDA) (Sigma-Aldrich), and 5 or 10 mM L-ascorbic acid (Sigma-Aldrich) final concentrations respectively. Control wells contained 500 µL per well of complete K-medium with 5 or 10 mM L-ascorbic acid. The 24-well plate was covered with foil and put on an orbital shaker at 20 °C and 120 RPM for one hour. Afterward, worms were collected in separate 15 mL conical tubes, the total volume raised to 15 mL with K-medium, and the worms were allowed to gravity-settle. After 3 minutes, the supernatant was discarded, and this washing step was repeated one more time. Microscopy was performed immediately after.

### 2.7. Statistical analysis

OASIS2 was used for analysis of paralysis induced by aldicarb and for analysis of reproduction per day [38]. MATLAB R2024b (MATLAB 24.2) was used for all other statistical testing and graph generation. Statistical tests and sample sizes are described in their corresponding figure legends.

## 3. Results

### 3.1. Inhibition of the Krebs cycle impaired growth but did not cause neurotoxicity

We aimed to test if developmental exposure to Krebs cycle inhibitors would cause dopaminergic neurotoxicity in *C. elegans*. To ensure that any neuronal damage we quantified was not the product of whole organism toxicity, we selected concentrations that caused mild growth delay, and we also evaluated neurotoxicity in glutamatergic and cholinergic neurons.

To impair function of the Krebs cycle we utilized sodium arsenite and fluoroacetic acid. Arsenic affects the Krebs cycle by inhibiting the enzymes pyruvate hydrogenase and isocitrate dehydrogenase [12, 39], while fluoroacetic acid is a precursor of fluorocitrate, an inhibitor of aconitase which is an enzyme in both the Krebs and glyoxylate cycles [23, 40]. Moreover, arsenic is a persistent pollutant in groundwater linked to increased cancer risk [2, 3], and fluoroacetate is a pesticide with the commercial name of Compound 1080 with impacts including livestock poisoning [41, 42]. Worms were exposed from the embryo stage for 52 to 54 hours until they reached the L4 larval stage, at which point all their somatic cells are fully formed (Figure 1A). All developmental endpoints were quantified at this point, including neuron morphological damage for all neuronal types tested (Figure 1B). We observed a significant decrease in normalized growth length for all sodium arsenite concentrations tested between 50 and 2000 µM (Figure 1C) and for all fluoroacetic acid concentrations tested between 750 and 1500 µM (Figure 1D). We found similar results for the non-normalized length values and for worm volume (<u>Figure A2</u>). We used this data to calculate concentrations causing a reduction of 1%, 10% and 25% in average worm length for sodium arsenite, corresponding to 0.1, 166.2, and 943.2 µM respectively. Similarly, we calculated the concentrations causing a reduction of 1%, 10% and 30% in average worm length for fluoroacetic acid to be 6.7, 75.2, and 390.0 µM respectively. We used these EC_1_, EC_10_ and EC_25_/EC_30_ values for subsequent experiments to enable us to both ensure that our neuronal endpoint measurements were not caused by generalized whole organism toxicity (EC_1_, EC_10_) and, in the case of negative results, ensure that our inability to detect an outcome was not the result of lack of uptake (EC_25_/EC_30_).

**Figure 1.**
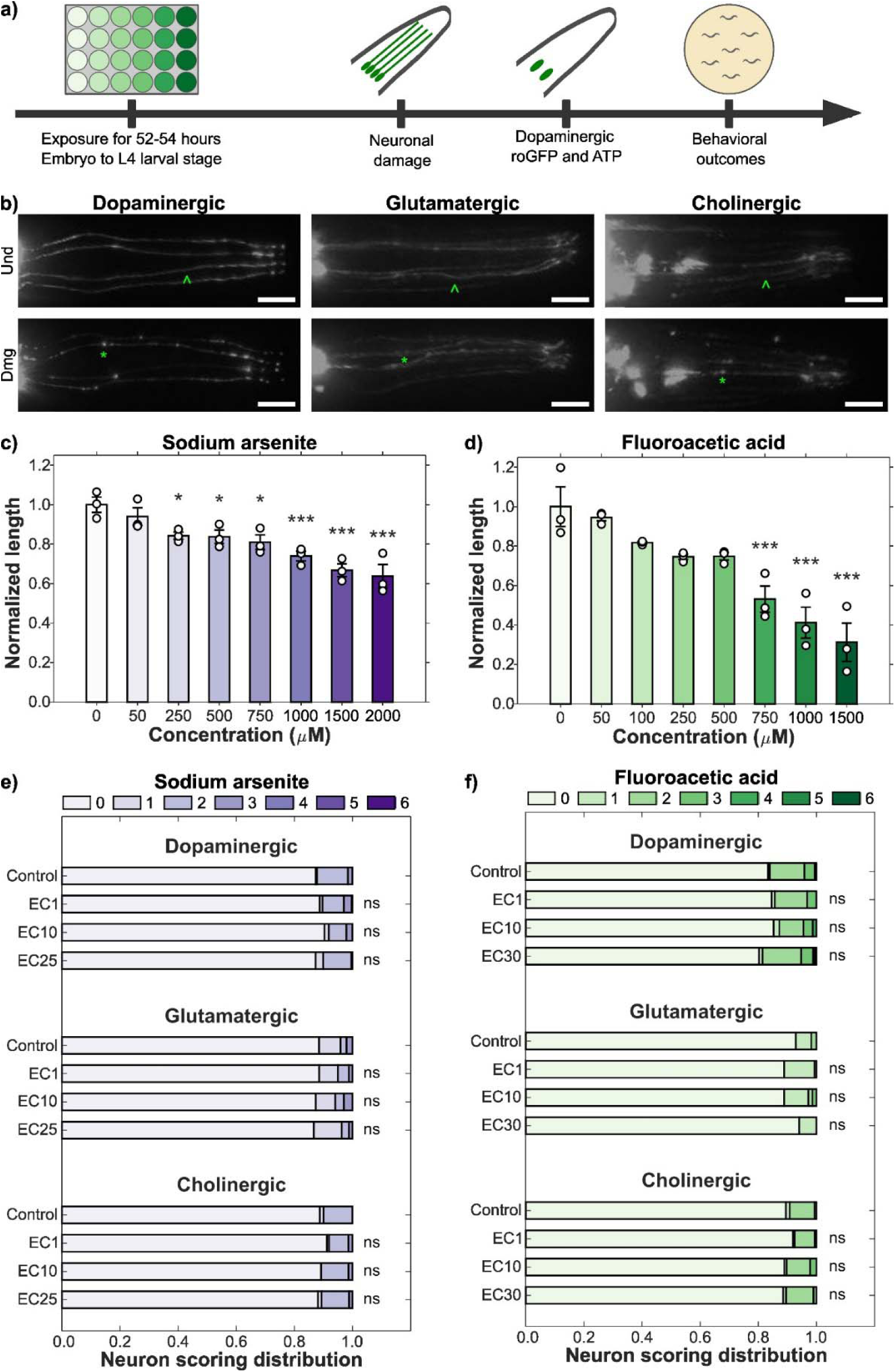
Krebs cycle inhibition does not cause developmental neurotoxicity. (a) Developmental exposure starts at the embryo stage until the L4 larval stage, when endpoint measurement is performed. (b) Representative images of undamaged (^) and damaged (*) neurites of cephalic dopaminergic, glutamatergic, and cholinergic neurons, scale bars 10 µm. Normalized length of worms exposed to increasing concentrations of (c) sodium arsenite and (d) fluoroacetic acid during development. *N* = 3 biological replicates, *n* = 50-100 worms per treatment, error bars SEM, one-way ANOVA with Dunnet’s post-hoc test, * *p* < 0.05, *** *p* < 0.001. Neuronal damage scoring distribution for dopaminergic, glutamatergic, and cholinergic neurons of worms developmentally exposed to (e) sodium arsenite and (f) fluoroacetic acid. *N* = 3 biological replicates, *n* = 70-80 dendrites per treatment, chi-square test with Bonferroni post-hoc test, ns *p* > 0.05.

We also tested if developmental exposure to sodium arsenite or fluoroacetic acid would cause reproductive impairment. We found that the total number of viable embryos from worms exposed to sodium arsenite significantly decreased for worms exposed to 1000 µM sodium arsenite and for worms exposed to 500 µM fluoroacetic acid (<u>Figure A3</u>). However, both 500 and 1000 µM sodium arsenite caused a significant reduction in reproduction (flattening of the curve when graphing the number of viable embryos per day). Fluoroacetic acid did not cause a significant decrease in viable embryos statistically assessed on a per-day basis at any concentration tested (<u>Figure A3</u>).

Next, we evaluated if these developmental exposures, causing mild and moderate growth inhibition, would cause neurotoxicity. We quantified neuron morphological damage in three distinct neuronal types (defined by neurotransmitter), quantifying morphological irregularities including kinks, blebs, and breaks in dendrites of neurons localized in the head of *C. elegans* (<u>Figure A1</u>) using previously described scoring systems [34, 35], with increasing scoring number corresponding with increased degrees of damage. Developmental exposure to sodium arsenite did not cause a significant neurotoxic effect in dopaminergic, glutamatergic, or cholinergic neuronal damage for any of the concentrations tested (Figure 1E). Similarly, developmental exposure to fluoroacetic acid did not cause a significant effect on any of the neuronal types evaluated (Figure 1F).

### 3.2. Inhibition of the Krebs cycle does not alter neuronal function

Beyond direct quantification of neuronal structural damage, impairment of neuronal function can be evaluated in *C. elegans* by measuring behaviors linked to specific neuron types [35]. For instance, we have previously shown that dopaminergic morphological damage in *C. elegans* is associated with a loss of their basal slowing response (BSR) behavior [43]; BSR is a food-scavenging behavior regulated by dopaminergic neurons in *C. elegans* [44]. Thus, we assessed whether Krebs cycle inhibition caused functional deficits that might appear in the absence of significant neuron morphological damage. In the BSR assay, control worms keep moving while searching for food, but move slower in the presence of bacterial food. This response was retained after developmental exposure to either sodium arsenite or fluoroacetic acid (Figure 2A). Mutant strains for *cat-2* and *cat-4*, included as positive controls, showed the expected loss of their basal slowing response.

**Figure 2.**
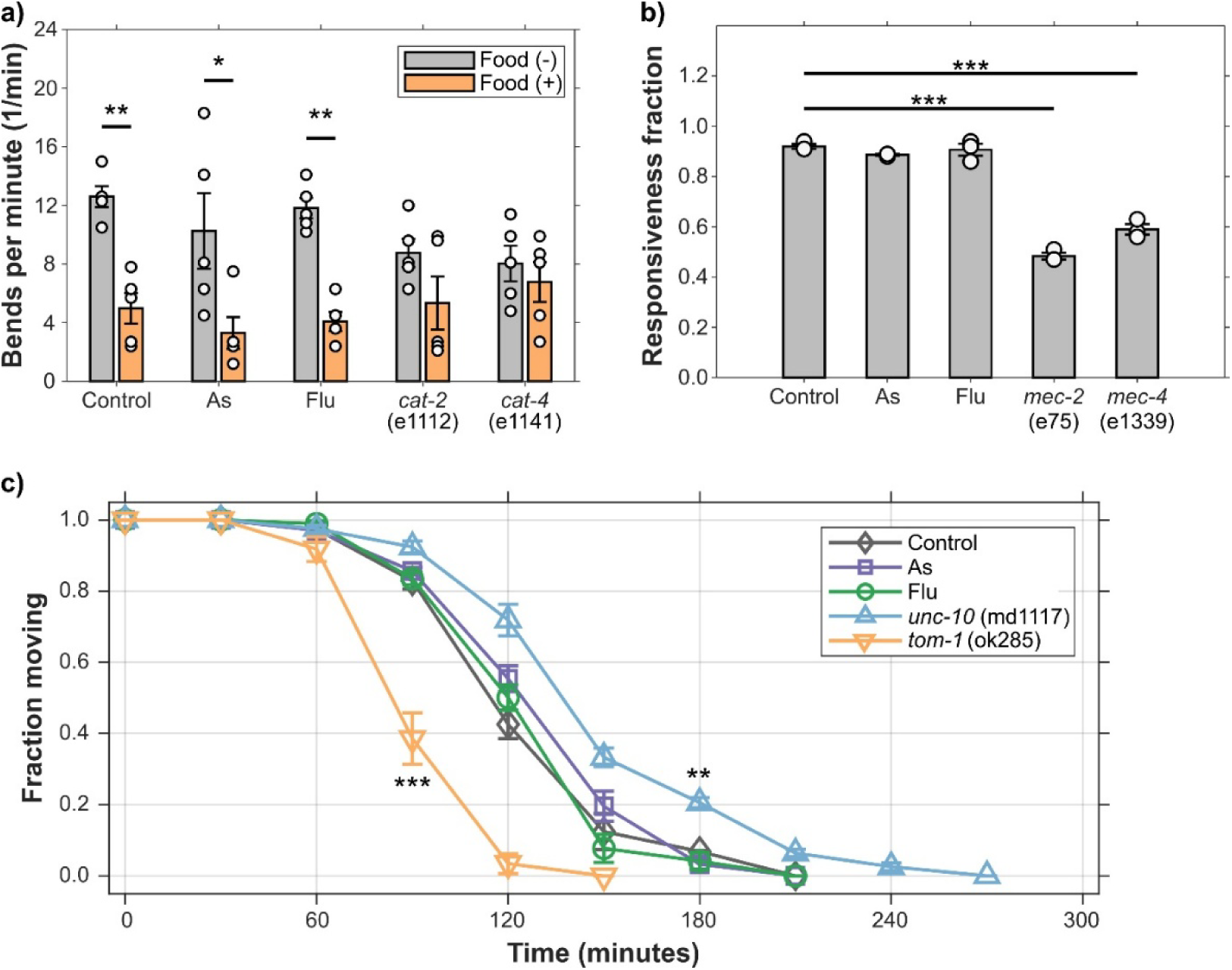
Krebs cycle inhibition does not alter behavior. (a) Basal slowing response, (b) sensitivity to soft touch, and (c) aldicarb-induced paralysis after developmental exposure to sodium arsenite and fluoroacetic acid. *N* = 5 biological replicates, *n* = 10 worms per treatment for (a), *N* = 3 biological replicates, *n* = 20 worms per treatment for (b), *N* = 3 biological replicates, *n* = 25-30 worms per treatment for (c), error bars SEM, two-way ANOVA with Tukey’s HSD post-hoc test for (a), one-way ANOVA with Dunnett’s post-hoc test for (b), log-rank test for (c), * *p* < 0.05, ** *p* < 0.01, *** *p* < 0.001.

We also evaluated the effects of developmental exposure on sensitivity to soft touch, a behavior regulated by glutamatergic mechanosensory neurons in the head and tail of worms [45]. After gentle prodding with an eyelash, control worms respond by recoiling from the source of the stimulus. Neither sodium arsenite nor fluoroacetic acid altered responsiveness to soft touch respect to control (Figure 2B). Strains with mutations in *mec-2* and *mec-4* were used as positive controls and showed significant reduction in responsiveness with respect to control.

Finally, we evaluated aldicarb-induced paralysis as an indicator of cholinergic function. Aldicarb is an insecticide that inhibits the enzyme acetylcholinesterase, disrupting the normal breakdown of acetylcholine [46]. Developmental exposure to sodium arsenite or fluoroacetic acid did not alter the rate of aldicarb-induced paralysis with respect to control (Figure 2C). Positive control worms with mutations in *unc-10* and *tom-1* showed resistance and hypersensitivity respectively. Together, these results demonstrate that developmental inhibition of the Krebs cycle alone does not produce detectable dopaminergic, glutamatergic, or cholinergic functional impairment at concentrations causing mild or moderate organismal toxicity.

### 3.3. Loss of mitophagy gene pink-1, but not pdr-1 or the presence of α-synuclein, enables dopaminergic neurotoxicity by sodium arsenite

We hypothesized that the absence of neurotoxicity after chemical exposure could reflect compensatory mechanisms that protect dopaminergic neurons from Krebs cycle disruption. Because Parkinson’s disease and other neurodegenerative diseases can arise through interactions between environmental exposures and genetic susceptibility [47, 48], we evaluated if impaired mitophagy by itself or in combination with dopaminergic expression of human α-synuclein [49] could sensitize worms to sodium arsenite or fluoroacetic acid.

Expression of α-synuclein alone caused substantial baseline morphological damage in dopaminergic neurons. However, neither sodium arsenite nor fluoroacetic acid further increased neuronal damage in the α-synuclein background for any of the concentrations we tested (Figure 3A-B). Similarly, loss of mitophagy gene *pdr-1*, the *C. elegans* homolog of human PRKN [50], did not increase dopaminergic neurotoxicity after exposure to either chemical (the apparent trend of sensitization to sodium arsenite was not statistically significant). However, sodium arsenite exposure at the EC_25_ concentration significantly increased dopaminergic neuronal damage in worms lacking *pink-1*, the *C. elegans* homolog of human PINK1 [51] (Figure 3A). This effect was also observed in worms carrying both dopaminergic α-synuclein and the *pink-1* mutation (Figure 3A). On the other hand, this effect of the loss of *pink-1* was not observed after fluoroacetic acid developmental exposure, or for any other of the tested genetic backgrounds (Figure 3B). Therefore, sodium arsenite, but not fluoroacetic acid, can produce dopaminergic neurotoxicity when mitochondrial quality-control capacity is compromised by loss of *pink-1*.

**Figure 3.**
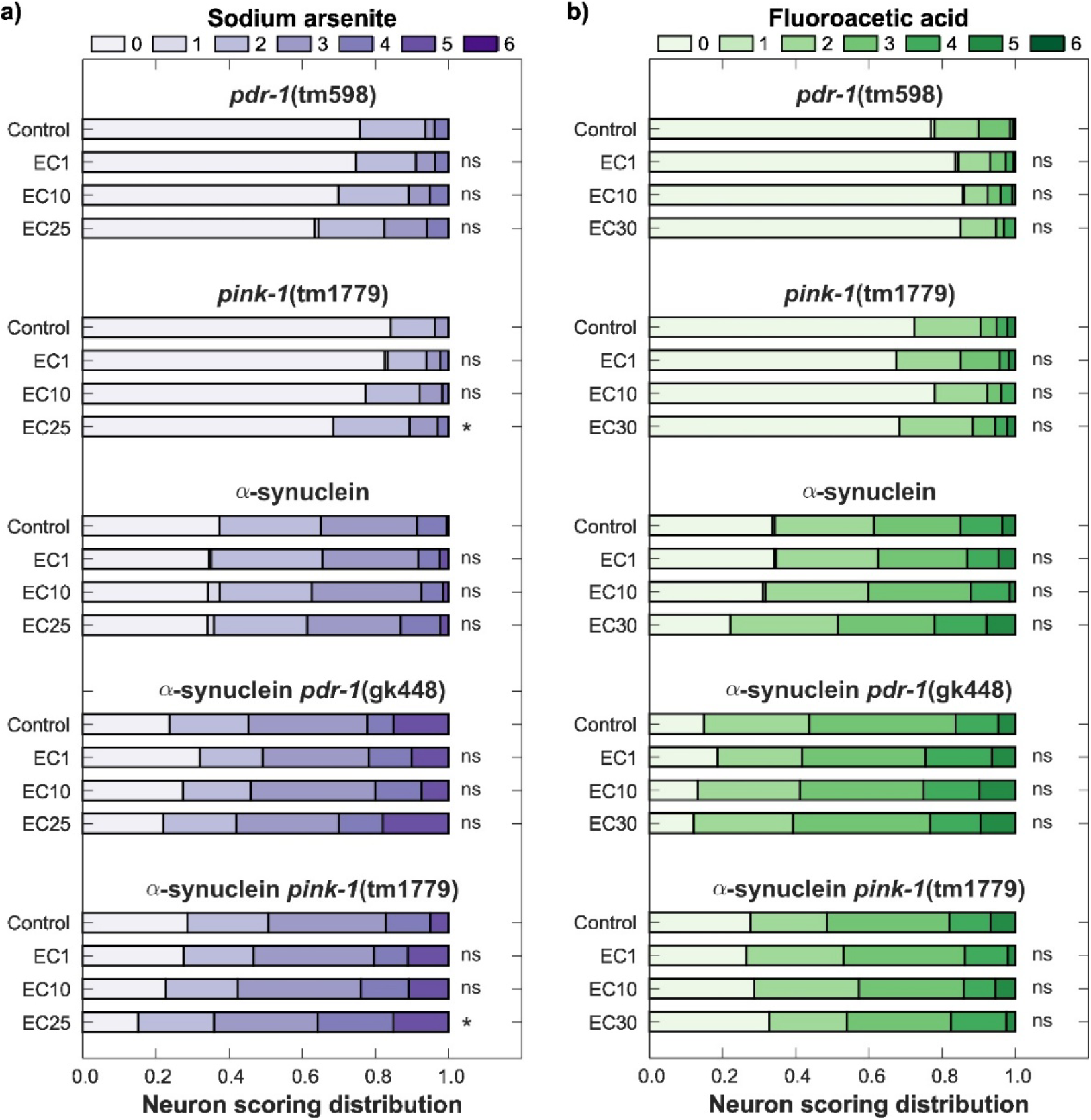
Sodium arsenite and loss of mitophagy gene *pink-1* together cause developmental dopaminergic neurotoxicity. Neuronal damage scoring distribution for cephalic dopaminergic neurons in combination with dopaminergic α-synuclein, a *pdr-1* mutant background, a *pink-1* mutant background, dopaminergic α-synuclein and *pdr-1* mutant background, and dopaminergic α-synuclein and *pink-1* mutant background in worms developmentally exposed (a) sodium arsenite and (b) fluoroacetic acid. *N* = 3 biological replicates, *n* = 70-80 dendrites per treatment, chi-square test with Bonferroni post-hoc test, ns *p* > 0.05, * *p* < 0.05.

### 3.4. Sodium arsenite reduces ATP levels in dopaminergic neurons

Having found that sodium arsenite can cause dopaminergic neurotoxicity in a mitophagy deficient background, we asked if the absence of neurotoxicity after chemical exposure alone might reflect limited effects on dopaminergic mitochondrial redox or bioenergetic states. To test this, we measured dopaminergic redox state in dopaminergic neurons with a mitochondrial roGFP reporter. This reduction-oxidation sensitive green fluorescent protein is a biosensor that captures variations in the glutathione pool (GSH/GSSG) [52]. Similarly, we used a dopaminergic PercevalHR reporter, that follows the same principles by reporting the ratio of ATP to ADP in dopaminergic neurons [53].

Neither sodium arsenite nor fluoroacetic acid significantly increased the dopaminergic mitochondrial roGFP ratio for any of the tested concentrations (Figure 4A-B). However, sodium arsenite significantly reduced the dopaminergic ATP:ADP ratio at both the EC_10_ and EC_25_ concentrations (Figure 4C). Fluoroacetic acid did not significantly decrease the dopaminergic ATP:ADP ratio (Figure 4D), although there was a non-significant trend towards decrease in the dopaminergic ATP:ADP ratio with increasing concentrations of this chemical. Hence, developmental exposure to sodium arsenite caused a significant dopaminergic bioenergetic deficit without a significant change in mitochondrial redox state as reported by roGFP. Conversely, fluoroacetic acid did not significantly alter either endpoint under these developmental exposure conditions. Because we observed a non-significant trend towards increasing dopaminergic roGFP ratio under sodium arsenite exposure, we also tested redox state with a whole-organism (ubiquitously expressed) cytosolic roGFP reporter. However, we found no effect of developmental exposure to sodium arsenite on whole body mitochondrial redox state as reported by roGFP (<u>Figure A4</u>). The non-significant trend to increased roGFP ratio observed in the dopaminergic neurons was not observed using this ubiquitous reporter.

**Figure 4.**
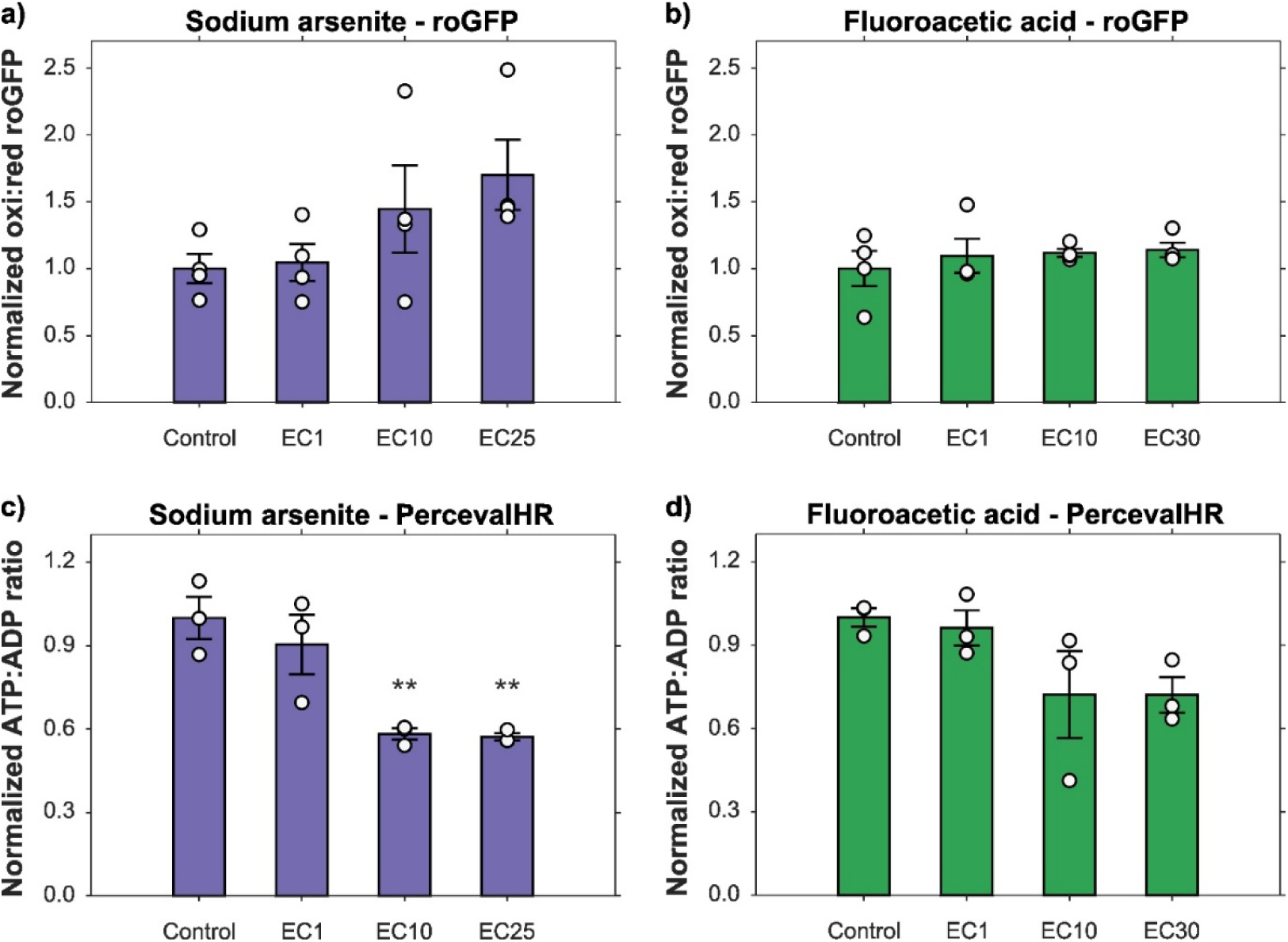
Sodium arsenite alters dopaminergic bioenergetic state. Oxidized to reduced roGFP ratio in cephalic dopaminergic neurons (cell bodies) of *C. elegans* developmentally exposed to (a) sodium arsenite and (b) fluoroacetic acid. *N* = 4 biological replicates, *n* = 40 neurons per treatment, error bars SEM, one-way ANOVA with Dunnett’s post-hoc test. ATP to ADP ratio in cephalic dopaminergic neurons (cell bodies) of *C. elegans* exposed (c) sodium arsenite and (d) fluoroacetic acid. *N* = 3 biological replicates, *n* = 40 neurons per treatment, error bars SEM, one-way ANOVA with Dunnett’s post-hoc test, ** *p* < 0.01. All roGFP and ATP to ADP values normalized to their respective controls.

### 3.5. Sensitization to redox stress enables sodium arsenite-induced dopaminergic neurotoxicity

The reduction in dopaminergic ATP:ADP ratio caused by sodium arsenite did not, by itself, result in neuronal morphological damage or loss of neuronal function. However, our previous work points to an important role of redox stress in developmental neurotoxicity [17, 18, 54]. Given the hint of altered redox state (Figure 4) and arsenic’s ability to bind sulfhydryl groups and cause redox stress [55, 56], we next tested whether chemical, genetic, or secondary toxicant challenges that increase oxidative stress or reduce mitochondrial resilience would also sensitize dopaminergic neurons to sodium arsenite.

First, we evaluated co-exposure to rotenone and antimycin A, inhibitors of mitochondrial electron transport chain complexes I and III respectively [57, 58]; both cause dopaminergic neuron redox stress in *C. elegans* [17, 18]. We also tested co-exposure to buthionine sulfoximine (BSO), which irreversibly inhibits γ-glutamylcysteine synthetase, the rate-limiting enzyme in glutathione synthesis, causing a sustained reduction in glutathione levels and increasing oxidative stress [59, 60]. Exposure to 1.0 µM rotenone, 0.1 µM antimycin A, or 500 µM buthionine sulfoximine alone did not significantly increase dopaminergic neuronal damage, and sodium arsenite EC_25_ alone also remained non-neurotoxic. However, co-exposure of sodium arsenite with 1.0 µM rotenone, 0.1 µM antimycin A, or 500 µM BSO significantly increased dopaminergic neuron morphological damage with respect to both control and sodium arsenite EC_25_ (Figure 5A). These results indicate that developmental exposure to sodium arsenite can produce dopaminergic neurotoxicity when combined with additional chemical disruption of mitochondrial function or glutathione-dependent antioxidant capacity.

**Figure 5.**
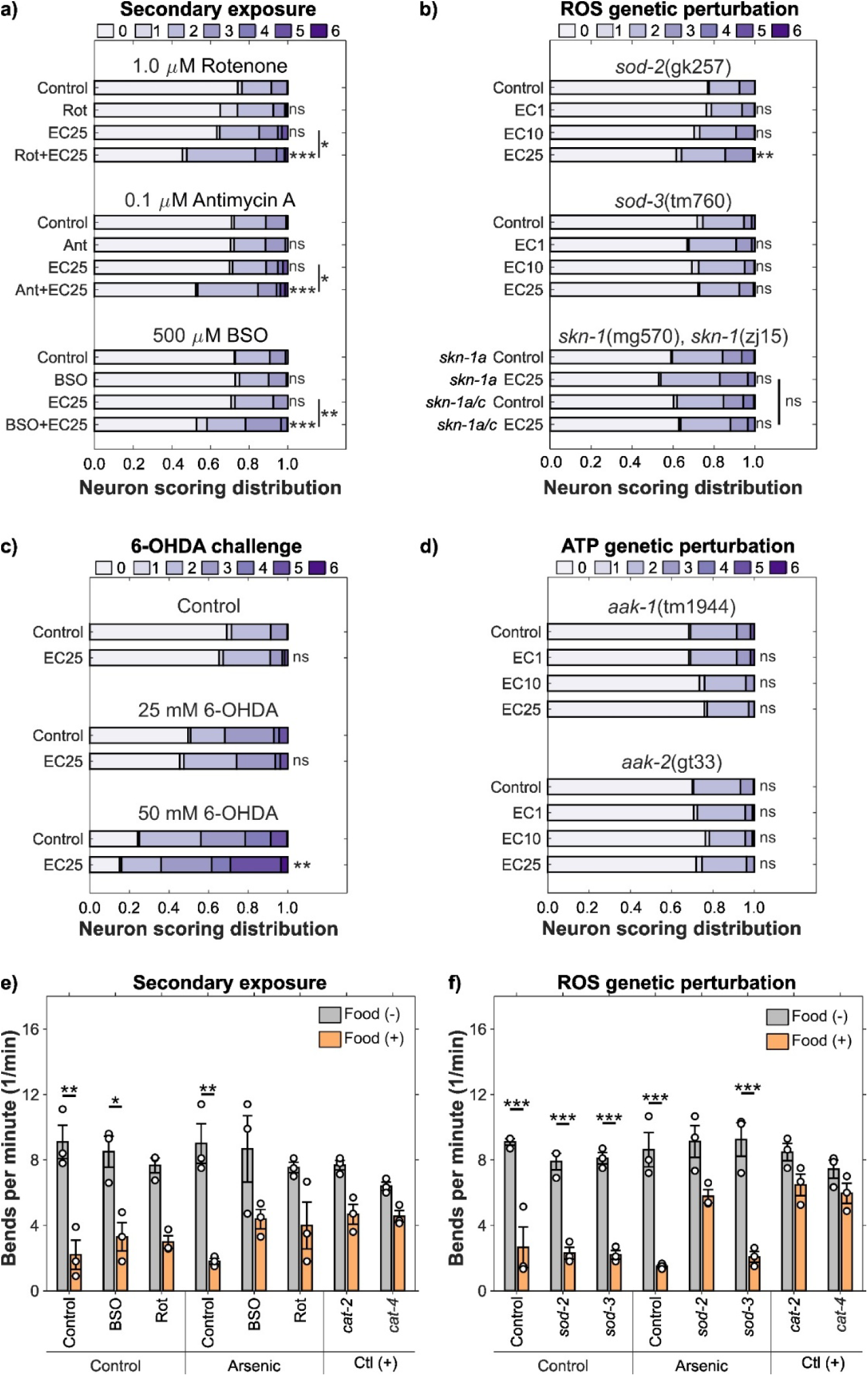
Sodium arsenite causes developmental dopaminergic neurotoxicity in the presence of additional redox stress. (a) Neuronal damage scoring distribution for cephalic dopaminergic neurons in combination with 1.0 µM rotenone, 0.1 µM antimycin A, and 500 µM buthionine sulfoximine. (b) Neuronal damage scoring distribution for cephalic dopaminergic neurons in *sod-2*, *sod-3*, and *skn-1* mutant backgrounds. (c) Neuronal damage scoring distribution for cephalic dopaminergic neurons after secondary challenge with 25 and 50 mM 6-OHDA. (d) Neuronal damage scoring distribution for cephalic dopaminergic neurons in *aak-1*, and *aak-2* mutant backgrounds. *N* = 3 biological replicates, *n* = 70-80 dendrites per treatment, chi-square test with Bonferroni post-hoc test, ns *p* > 0.05, * *p* < 0.05, ** *p* < 0.01, *** *p* < 0.001. (e) Basal slowing response for worms exposed to sodium arsenite EC_25_ in combination with 500 µM buthionine sulfoximine, and 1.0 µM rotenone. (f) Basal slowing response for worms exposed to sodium arsenite EC_25_ in *sod-2*, and *sod-3* mutant backgrounds. *N* = 3 biological replicates, *n* = 10 worms per treatment, error bars SEM, three-way ANOVA with Tukey’s HSD post-hoc test, * *p* < 0.05, ** *p* < 0.01, *** *p* < 0.001.

The oxidative stress response of *C. elegans* can be also disrupted by genetic intervention. Thus, we evaluated the effects of developmental exposure to sodium arsenite in combination with loss of the mitochondrial superoxide dismutase *sod-2* and *sod-3* genes. Deletion of *sod-2* increases sensitivity to oxidative stress by reducing turnover of superoxide radicals into hydrogen peroxide [61]; as does loss of *sod-3* [62], although the expression pattern of these enzymes varies from each other [63, 64]. We also tested two isoform-specific mutants of *skn-1*, the *C. elegan*s ortholog of Nrf2 [65]. The isoform *skn-1a* regulates proteasome capacity and protein quality control, while *skn-1a/c* also activates the oxidative stress response [66, 67]. Loss of *sod-2* enabled significant dopaminergic neurotoxicity after developmental exposure to sodium arsenite EC_25_ (Figure 5B). On the other hand, loss of *sod-3* did not lead to sensitization to dopaminergic neurotoxicity after sodium arsenite exposure. Similarly, neither *skn-1a* nor *skn-1a/c* mutant backgrounds increased dopaminergic neurotoxicity after sodium arsenite exposure (Figure 5B). These results identify loss of *sod-2*, but not the other redox-response mutations tested, as a genetic condition that permits sodium arsenite neurotoxicity in dopamine neurons.

Next, we evaluated if developmental exposure to sodium arsenite sensitized dopaminergic neurons to a secondary challenge with the well-characterized dopaminergic neurotoxin 6-hydroxydopamine (6-OHDA) [68]. Sodium arsenite EC_25_ by itself did not increase neuronal damage, and it did not enhance damage caused by 25 mM 6-OHDA. However, worms developmentally exposed to sodium arsenite EC_25_ showed significantly increased dopaminergic neuron morphological damage after challenge with 50 mM 6-OHDA compared to worms exposed only to 50 mM 6-OHDA (Figure 5C). Thus, developmental sodium arsenite exposure can increase susceptibility to a subsequent dopaminergic toxicant challenge.

Because sodium arsenite EC_10_ and EC_25_ significantly reduced the ATP:ADP ratio in dopaminergic neurons, we also tested whether impaired AMP-activated protein kinase signaling sensitized neurons to sodium arsenite. This genetic pathway acts as energy sensor that activates when ATP levels drop and the AMP:ATP ratio increases [69, 70]. Loss of either of the two genes encoding the catalytic subunits of AMP-activated protein kinase, *aak-1* and *aak-2*, did not cause dopaminergic neurotoxicity after developmental exposure to sodium arsenite (Figure 5D), indicating that loss of these energy-sensing pathways is not sufficient to sensitize dopamine neurons to potential sodium arsenite-linked neurotoxicity.

Finally, we tested whether the neuron morphological effects observed after redox-stress sensitization were accompanied by impaired dopaminergic function as measured by BSR. Developmental exposure to sodium arsenite EC_25_ combined with 500 µM BSO or 1.0 µM rotenone abrogated the food-slowing behavior of the basal slowing response relative to the corresponding rate of movement on plates without food (Figure 5E). Notably, this loss of the basal slowing response was also observed for the single chemical exposure to 1.0 µM rotenone, a complex I inhibitor previously reported to impair dopaminergic neuronal function in *C. elegans* [18]. Furthermore, sodium arsenite EC_25_ exposure in the *sod-2* mutant background also impaired the basal slowing response, although this behavioral effect was not observed in the *sod-3* background (Figure 5F). We also tested mitochondrial redox state in dopaminergic neurons after combinatorial exposure to sodium arsenite EC_10_ and EC_25_ with 1.0 µM rotenone and 500 µM BSO using the roGFP reporter. We found a significant increase in oxidized to reduced dopaminergic roGFP ratio for sodium arsenite EC_10_ and EC_25_ by themselves, and for their combinations with rotenone and BSO (<u>Figure A5</u>). We did not find a significant increase in oxidized to reduced roGFP for the co-exposures with respect to their single-chemical sodium arsenite results. It is plausible that sodium arsenite causes a response from the roGFP reporter under these conditions such that the fluorescence intensity reaches a maximum, prohibiting measurement of larger responses. These results show that sodium arsenite produces both morphological damage and functional dopaminergic deficits when redox stress response is compromised.

## 4. Discussion

Environmental pollutants can disrupt mitochondrial function through several pathways, and mitochondrial dysfunction is associated with neurotoxicity, especially of the dopamine neurons. However, only a few specific modes of mitochondrial dysfunction have been tested for selective dopaminergic toxicity. In this work, we evaluated whether developmental inhibition of the Krebs cycle would cause dopaminergic neurotoxicity *in vivo* in *C. elegans* using sodium arsenite and fluoroacetic acid. Both Krebs cycle inhibitors impaired growth and reproduction, but neither caused either morphological damage or functional loss in dopaminergic, glutamatergic, or cholinergic neurons at concentrations causing mild or moderate organismal toxicity. However, sodium arsenite exposure caused dopaminergic neuron morphological damage and functional deficit when redox or mitochondrial stress response capacity was reduced chemically, genetically, or by secondary chemical challenge. Thus, developmental exposure to sodium arsenite requires additional redox stress to induce dopaminergic neurotoxicity. However, our results do not support the hypothesis that Krebs cycle disruption during development is by itself sufficient to cause dopaminergic neurotoxicity in *C. elegans*, unlike inhibition of mitochondrial complexes I and III.

It is important to note that sodium arsenite and fluoroacetic acid are not equivalent in their mode of action or selectivity in targeting the Krebs cycle. Sodium arsenite has multiple targets, including enzymes such as pyruvate dehydrogenase, while fluoroacetic acid is converted to fluorocitrate which inhibits aconitase. Previous research in *C. elegans* has found that arsenite decreases pyruvate dehydrogenase activity, ATP-linked respiration, and spare respiratory capacity while increasing proton leak and glycolytic reliance at the whole-organism level [12, 13]. Our results showing significantly reduced dopaminergic ATP levels are consistent with these established bioenergetic effects of arsenic in *C. elegans*, while extending this effect to a specific neuronal type. Fluoroacetic acid is generally considered to be a selective inhibitor (after metabolism to fluorocitrate) of aconitase [71, 72]. We did not find dopaminergic neurotoxicity after developmental exposure to fluoroacetic acid, despite observing significant effects on growth and reproduction. A previous report found strong and relatively reproduction-selective effects of fluoroacetic acid in *C. elegans* [23]. However, their findings also challenged the general idea that fluoroacetic acid acts specifically by targeting aconitase. Combined inhibition of the Krebs and glyoxylate cycles, with malonate and itaconate respectively, did not recapitulate the effects of fluoroacetic acid on reproduction, suggesting that fluoroacetic acid exposure outcomes may not come only from aconitase inhibition. Nonetheless, regardless of whether arsenite and fluoroacetate have effects outside the Krebs cycle, there is no question that both do inhibit the Krebs cycle. Thus, our results do not support that Krebs cycle inhibition alone leads to degeneration of dopaminergic, glutamatergic, or cholinergic neurons at concentrations causing a 25-30% reduction in growth.

Our dopaminergic neurotoxic effects are dissimilar to those previously reported of impaired locomotor behavior and dopaminergic neuronal degeneration after continuous developmental exposure to 1 and 5 mM sodium arsenite in *C. elegans*, which was accompanied by induction of antioxidant genes *skn-1*, *gcs-1*, and *gss-1* [73]. It is possible that we would have seen dopaminergic neurodegeneration at higher exposure concentrations, since our highest tested concentration (943.2 µM) was slightly lower than the lowest concentration that caused a detectable effect in this previous work. Alternatively, several exposure parameters were different between our studies, including culture medium, developmental timing, duration of exposure, reporter strain, and neuronal scoring method, which could have influenced the outcome.

Similarly, studies in vertebrate models show that arsenic exposure can lead to dopaminergic neurodegeneration, but the outcome varies between models, concentrations, developmental stage, and endpoints measured. In mice, 28-day arsenic exposure caused loss of nigrostriatal dopaminergic neurons, motor deficits, and inhibition of mitochondrial complexes II and IV [74]; while lower-level arsenic exposure increased α-synuclein accumulation in the brain and caused proteotoxic stress [75]. In zebrafish, chronic dietary arsenic exposure impaired cognition, oxidative stress, and altered dopaminergic signaling [76], while sodium arsenite exposure in embryos affected development of tyrosine hydroxylase-positive dopaminergic neurons [77].

Our results with dopaminergic redox and bioenergetic reporters show that ATP deficit in these neurons is possible without a significant effect on dopaminergic neurotoxicity. This is consistent with a study using the mitochondrial uncoupler pentachlorophenol, which caused decreased ATP and spare respiratory capacity after exposure without causing dopaminergic morphological damage or enhancing dopaminergic neurotoxicity of 6-OHDA [54]. Moreover, they did not observe a more-oxidized redox state, which is like our results with sodium arsenite alone.

Therefore, loss of bioenergetic capacity remains an important sign of mitochondrial stress under our exposure paradigm but is not, by itself, a reliable predictor of dopaminergic neurotoxicity.

On the other hand, a previous report using a rat model showed that chronic exposure to fluorocitrate causing astrocyte death did not cause dopaminergic neurodegeneration by itself but accelerated dopaminergic neuron damage from 6-OHDA [78]. This points to the importance of considering additional chemical sensitization to induce neurotoxicity. Consistent with this, our sensitization experiments show that sodium arsenite caused dopaminergic morphological damage and loss of basal slowing response when combined with the mitochondrial toxicants rotenone and antimycin A, the glutathione-depleting chemical BSO, and genetic loss of the mitochondrial superoxide dismutase gene *sod-2*. In comparison, loss of the genes encoding the AMPK subunits *aak-1* and *aak-2* did not enable dopaminergic neurotoxicity despite the already reduced dopaminergic ATP levels. These results confirm that ATP depletion is not the main driver of dopaminergic neurotoxicity by itself. Instead, it suggests the possibility that impaired bioenergetics may lower the threshold at which redox stress becomes damaging. Interestingly, sensitization to dopaminergic neurotoxicity was caused by loss of *sod-2*, but not *sod-3*, despite both mitochondrial superoxide dismutases having nonredundant interactions with electron transport chain function [79]. However, *sod-2* appears to be the major constitutively expressed mitochondrial SOD protein in *C. elegans* [80, 81], indicating that differences in the spatiotemporal dynamics between *sod-2* and *sod-3* could explain their distinct degrees of sensitization to neurotoxicity. *pink-1* and *pdr-1* are the *C. elegans* homologs of the Parkinson’s disease genes PINK1 and PRKN, and strains with mutations in these genes exhibit altered mitochondrial respiration and morphology [82]. *pink-1* and *pdr-1* were sensitizers for many chemicals in larval-growth assays [13], but this was interestingly not the case for arsenite. Thus, our identification of *pink-1* as a sensitizing mutant background for arsenic-mediated dopaminergic neurotoxicity indicates a selective sensitization of neurons. Another study found that *pdr-1* and *pink-1* mutations sensitized to dopaminergic neurodegeneration following ultraviolet C radiation, but *pdr-1* sensitized while *pink-1* protected against 6-OHDA-mediated dopaminergic neurotoxicity [83]. *pdr-1* and *pink-1* mutations also influence the dopaminergic neurotoxicity of manganese [84] and a neurotoxic bacterial metabolite [85]. In mice, a rotenone treatment that did not produce overt mitochondrial deficit impaired dopamine-dependent corticostriatal synaptic plasticity in Pink1 heterozygous animals [86], and intestinal infection induced Parkinson’s disease-like motor impairment selectively in Pink1 deficient animals [87]. However, another study found that mice with a Prkn defect did not show increased susceptibility to developmental manganese exposure [88]. Our work provides experimental support for gene–environment and environment-environment (chemical mixture) susceptibility, with arsenite-linked mitochondrial deficit becoming neurotoxic only when mitochondrial quality control (*pink-1*/PINK1) or antioxidant defenses (*sod-2*/SOD2) are compromised, consistent with prior evidence linking mitochondrial homeostasis genes and environmental stressors in *C. elegans* models of Parkinson’s disease [48]. Overall, it appears that gene-environment interactions depend on toxicological endpoint and that vulnerability to neurotoxicity cannot be inferred only from whole-organism measures.

The lack of evidence for Krebs cycle inhibition causing dopaminergic neurotoxicity that we report adds to other evidence [17, 18, 54, 83, 89] that mitochondrial dysfunction should not be treated as a single mode of action, but as distinct pathways with distinct neurotoxic outcomes. Developmental inhibition of mitochondrial complexes I and III was sufficient to cause selective dopaminergic neurotoxicity through site-specific mitochondrial reactive oxygen species production [17, 18]. In contrast, in this study, sodium arsenite required an additional deleterious redox stressor to elicit dopaminergic neurotoxicity. Further bioenergetic exacerbation via *aak-1* or *aak-2* mutation did not elicit dopaminergic neurodegeneration. This result contrasts with arguments that the sensitivity of dopamine neurons to mitochondrial dysfunction is attributable to their high energy demand [90, 91]. An alternate hypothesis is that the high energy reliance of dopaminergic neurons is important specifically in the context of combined stressors; i.e., energetic deficiency may sensitize to redox stress or vice versa. This points to the importance of studying environmental pollutants that could interact with this vulnerability to enact neurotoxicity, such as arsenic, a pervasive chemical in all aspects of human activity [92].

## Data availability

Data is publicly available at Duke Research Data Repository via the DOI: https://doi.org/10.7924/r4r589

## Author contributions

Conceptualization J.H. and J.N.M.; Method Development J.H. and J.N.M.; Formal Analysis J.H., W.E.T., and J.J.; Writing: Original Draft Preparation J.H. and J.N.M.; Writing: Review and Editing J.H. and J.N.M.; Supervision J.N.M.; and Funding Acquisition J.N.M.

