## Supplemental data for "Sodium arsenite requires redox sensitization to cause developmental dopaminergic neurotoxicity in *Caenorhabditis elegans*"

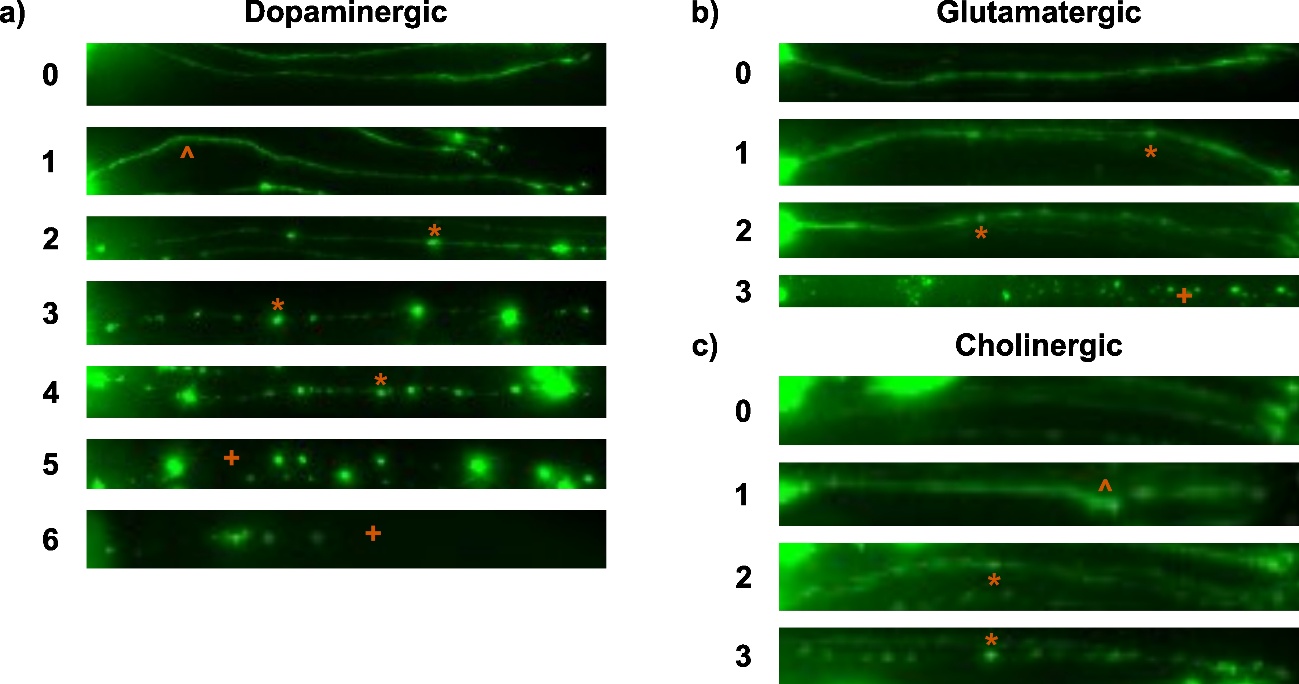

**Figure A1. Example images for neuronal damage scoring.** Dendrite damage for (a) dopaminergic neurons on a 0 to 6 scale, (b) glutamatergic neurons in a 0 to 3 scale, and (c) cholinergic neurons in a 0 to 3 scale. Typical types of damage include kinks (^), blebs (*), and breaks (+).

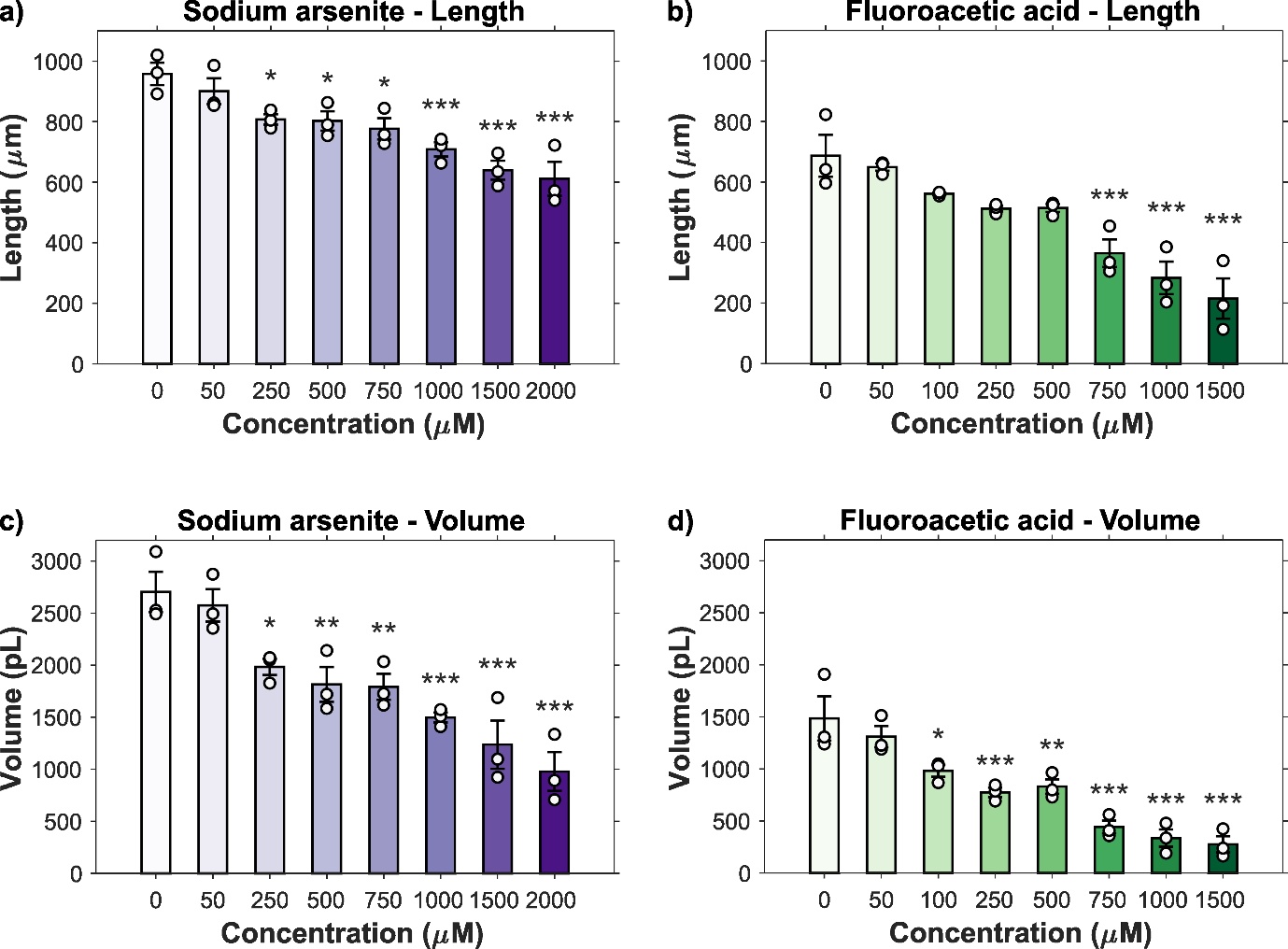

**Figure A2. Additional growth metrics.** Length of worms exposed to increasing concentrations of (a) sodium arsenite and (b) fluoroacetic acid during development. Volume of worms exposed to increasing concentrations of (c) sodium arsenite and (d) fluoroacetic acid during development. *N* = 3 biological replicates, *n* = 50-100 worms per treatment, error bars SEM, one-way ANOVA with Dunnet’s post-hoc test, * *p* < 0.05, ** *p* < 0.01, *** *p* < 0.001.

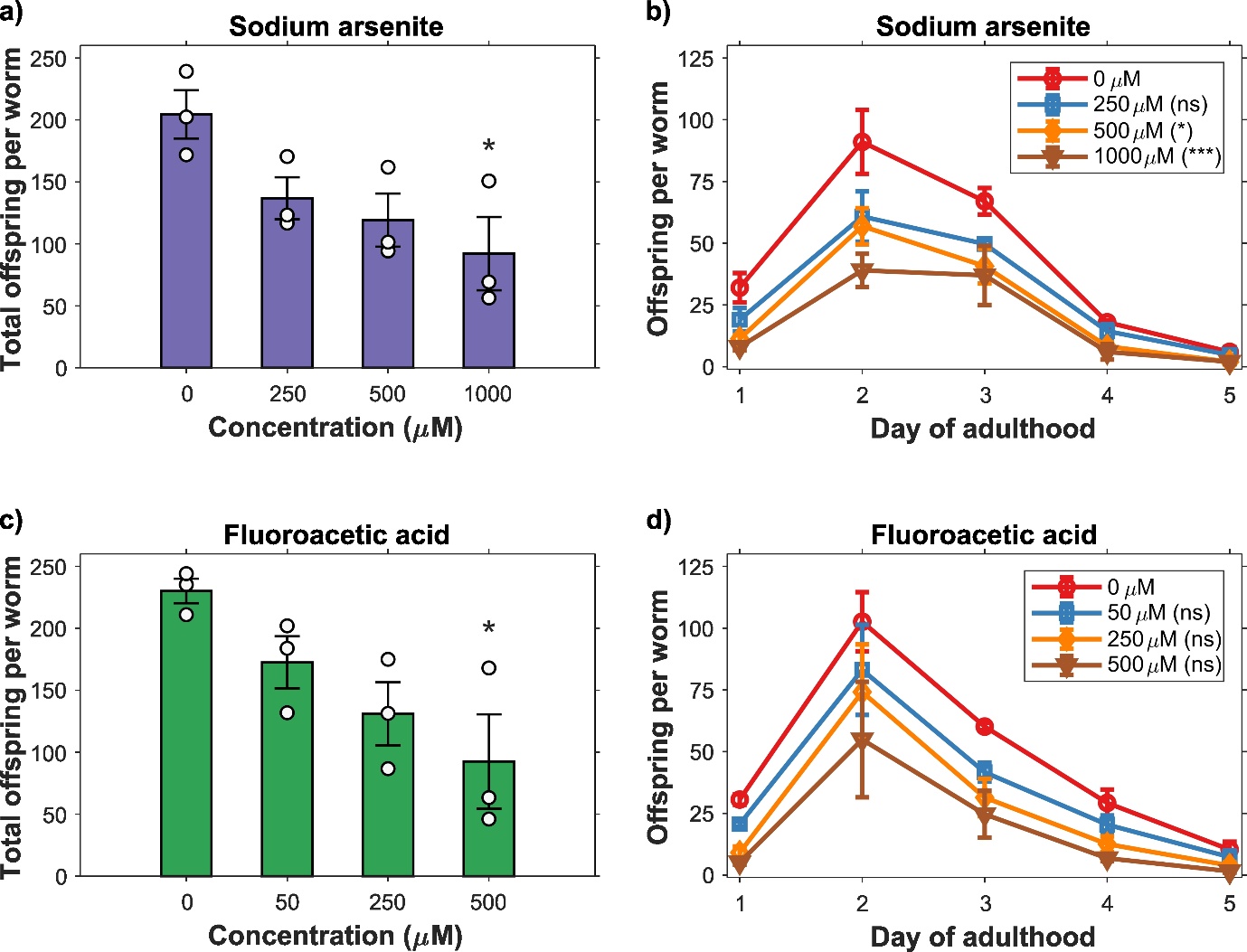

**Figure A3. Effects of developmental exposure on reproduction.** (a) Total viable offspring and (b) viable offspring per day of worms developmentally exposed to increasing concentrations of sodium arsenite. (c) Total viable offspring and (d) viable offspring per day of worms exposed during development to increasing concentrations of fluoroacetic acid. *N* = 3 biological replicates, *n* = 8-10 worms per treatment, error bars SEM, one-way ANOVA with Dunnet’s post-hoc test for (a) and (c), log-rank test for (b) and (d), ns *p* >0.05, * *p* < 0.05, *** *p* < 0.001.

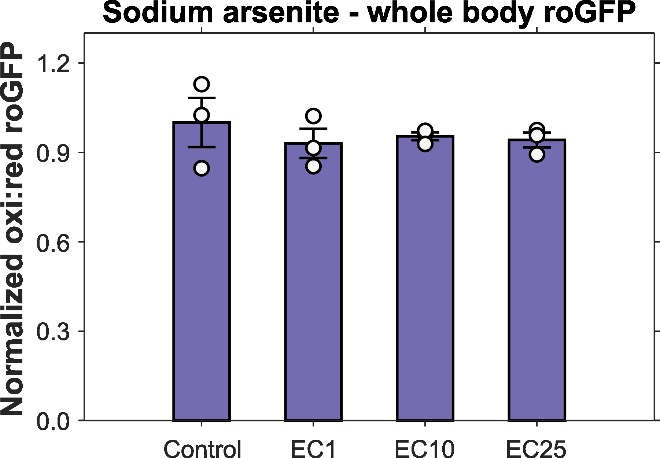

**Figure A4. Whole-body redox state after sodium arsenite exposure.** Oxidized to reduced cellular roGFP ratio in the whole body of *C. elegans* (ubiquitous reporter expression) developmentally exposed to sodium arsenite. *N* = 3 biological replicates, *n* = 20 worms per treatment, error bars SEM, one-way ANOVA with Dunnet’s post-hoc test. All roGFP ratio values normalized to their respective controls.

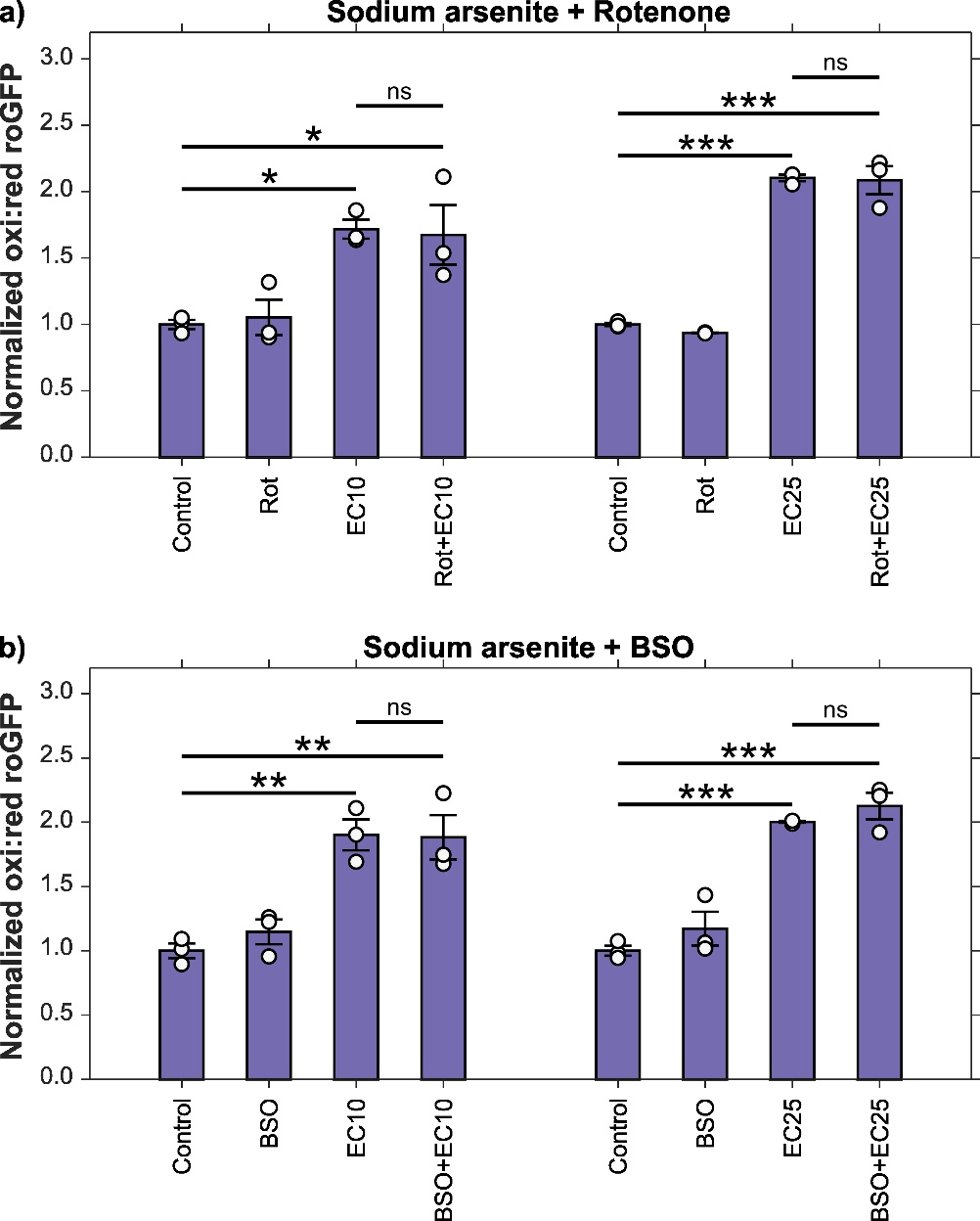

**Figure A5. Dopaminergic redox state after sensitization.** Oxidized to reduced roGFP ratio in cephalic dopaminergic neurons of *C. elegans* developmentally exposed to sodium arsenite EC_10_ and EC_25_ in combination with (a) 1.0 µM rotenone and (b) 500 µM buthionine sulfoximine. *N* = 3 biological replicates, *n* = 40 neurons per treatment, error bars SEM, two-way ANOVA with Tukey’s HSD post-hoc test for each treatment combination separately, ns *p* > 0.05, * *p* < 0.05, ** *p* < 0.01, *** *p* < 0.001. All roGFP ratio values normalized to their respective controls.

**Table A1. *C. elegans* strains.**

| **Strain** | **Genotype** | **Description** |
| --- | --- | --- |
| BY200 | *vtIs1*[p*dat-1*::GFP] | Dopaminergic reporter |
| DA1240 | *adIs1240*[*eat-4*::sGFP+*lin-15*(+)] | Glutamatergic reporter |
| LX929 | *vsIs48*[*unc-17*::GFP] | Cholinergic reporter |
| CB1112 | *cat-2*(e1112) II | BSR defective strain |
| CB1141 | *cat-4*(e1141) V | BSR defective strain |
| CB75 | *mec-2*(e75) X | Soft touch defective strain |
| CB1339 | *mec-4*(e1339) X | Soft touch defective strain |
| NM1657 | *unc-10*(md1117) X | Aldicarb resistant strain |
| VC223 | *tom-1*(ok285) I | Aldicarb sensitive strain |
| UA226 | *pink-1*(tm1779); *vtIs1*[p*dat-1*::GFP] | Dopaminergic reporter, *pink-1* mutant |
| UA227 | *pdr-1*(tm598); *vtIs1*[p*dat-1*::GFP] | Dopaminergic reporter, *pdr-1* mutant |
| ERS1 | *eraIs1*[*dat-1p*::mCherry, *dat-1p*::hSNCA::Venus] | Dopaminergic α-synuclein |
| ERS44 | *eraIs1*[*dat-1p*::mCherry, *dat-1p*::hSNCA::Venus]; *pdr-1*(gk448) | Dopaminergic α-synuclein, *pdr-1* mutant |
| ERS49 | *eraIs1*[*dat-1p*::mCherry, *dat-1p*::hSNCA::Venus]; *pink-1(tm1779)* | Dopaminergic α-synuclein, *pink-1* mutant |
| PHX2867 | p*dat-1*::MLS::roGFP | Dopaminergic roGFP reporter |
| PHX2923 | p*dat-1*::PercevalHR | Dopaminergic PercevalHR reporter |
| JV2 | *jrIs2*[*rpl-17*p::Grx1-roGFP2 + *unc-119*(+)] | Ubiquitous roGFP reporter |
| BY200 x GA184 | *vtIs1*[p*dat-1*::GFP]; *sod-2*(gk257) I | Dopaminergic reporter, *sod-2* mutant |
| BY200 x GA186 | *vtIs1*[p*dat-1*::GFP]; *sod-3*(tm760) X | Dopaminergic reporter, *sod-3* mutant |
| BY200 x GR2245 | *vtIs1*[p*dat-1*::GFP]; *skn-1*(mg570) IV | Dopaminergic reporter, *skn-1a* mutant |
| BY200 x QV225 | *vtIs1*[p*dat-1*::GFP]; *skn-1*(zj15) IV | Dopaminergic reporter, *skn-1a/c* mutant |
| BY200 x tm1944 | *vtIs1*[p*dat-1*::GFP]; *aak-1*(tm1944) III | Dopaminergic reporter, *aak-1* mutant |
| BY200 x TG38 | *vtIs1*[p*dat-1*::GFP]; *aak-2*(gt33) X | Dopaminergic reporter, *aak-2* mutant |

**Table A2. Primers used for PCR genotyping.**

| **Strain** | **Forward primer** | **Reverse primer** |
| --- | --- | --- |
| GR2245 (set 1) | AGTCGATGGGCGGTTCATCA | CACGTAACAATGCGGGAGAC |
| GR2245 (set 2) | CAGTCAGTCGATGGGCGGTT | ACTCGACTCTCAATTGTGGTGTAT |
| tm1944 | ACATCTTGATCAAATTAGCACTCTT | ATCAACTTCAGGGCCCACAT |
| TG38 | TTCATTTGCTGCAACTTCCTGG | ATGTTGTCGTAATCCTTTGCCA |
